# PSF correction in soft x-ray tomography

**DOI:** 10.1101/260737

**Authors:** Axel Ekman, Venera Weinhardt, Jian-Hua Chen, Gerry McDermott, Mark A. Le Gros, Carolyn Larabell

## Abstract

In this manuscript, we introduce a linear approximation of the forward model of soft x-ray tomography (SXT), such that the reconstruction is solvable by standard iterative schemes. This linear model takes into account the three-dimensional point spread function (PSF) of the optical system, which consequently enhances the reconstruction data. The feasibility of the model is demonstrated on both simulated and experimental data, based on theoretically estimated and experimentally measured PSFs.

## 1. Introduction

Soft x-ray tomography (SXT) refers to the x-ray microscopy technique in which tomographic imaging is done using low-energy x-rays. In particular, the x-ray energy range lies within the “water window”, i.e., between the K-absorption edges of oxygen (2.34 nm; 530 eV) and carbon (4.4 nm; 280 eV) [1]. As the name suggests, water is relatively transparent to the x-rays within this region. In biological samples, the contrast comes from the natural variation of bio-organic molecules making this region especially suitable for imaging of these kinds of samples. In the past decades, soft x-ray microscopy has emerged as a unique tool to study 3D organization of single cells [3, 6, 28]. From simple yeast to complex eukaryotic cells. SXT grants an unprecedented contrast and spatial resolution, filling the information gap between light and electron microscopy [33, 30, 10, 5, 17, 4].

The three-dimensional reconstruction of a sample is obtained, by solving for its spatial distribution of absorption from a series of projection from many different viewing angles around a central rotation axis. A key assumption has traditionally been that these images can be regarded as images of classical projections [37], meaning that the resolution would be limited by that of the optical system, *r* ∝ λ/*NA*, where λ is the wavelength of the illuminating light and *NA* is the numerical aperture of the objective lens. Diffraction limited optics are, however, also characterized by a maximum depth of field as DOF *∝* λ/*NA*^2^, introducing a limit of sample size where the image formation can be approximated by this kind of parallel projection [24], and where the effect of the optics can be modeled by a depth independent point spread function (PSF).

In principle, both a sufficient depth of field and a good resolution can be obtained by increasing the energy (decreasing λ) and decreasing the *NA* (see Fig. 1). However, natural contrast in bio-organic samples is limited to the energy region of the water window, making such optimization impossible. This means that, especially for larger samples, SXT resolution suffers from the depth-dependent optical PSF.

**Figure 1:**
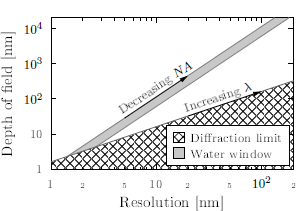
The relation between resolution and the depth of field for an ideal lens and monochromatic light. The diffraction limit shown here is defined as *r* = 0.61λ/*NA* for *NA* = 1. The water window shows accessible resolution and depth of field for soft x-ray energy range.

Limited DOF is a known problem in electron microscopy, where tilting of the sample may extend part of the sample out of focus, and approximate solutions exist to correct for it in the tomographic reconstruction. The so-called defocus-gradient correction, which was first introduced by Jensen and Kornberg [11], involves computational correction of the effects of a depth-dependent defocus and was incorporated to the well-known back-projection algorithm. The problem was revisited by Kazantsev et al. [13], in which the method received rigorous mathematical justification. This kind of correction has also been applied in Fourier space [36, 35], which can reduce the computational times up to two orders of magnitude. However, these corrections are not directly applicable in SXT due to the difference in image formation [16].

In SXT the problem of limited DOF has only recently been addressed, and thus far only experimentally. This has been done by acquiring multiple images at different focus and using a depth-dependent weight on their back-projection to account for their different foci [32], by using through-focus imaging to then computationally extract the ideal projection [22] or by a wavelet based fusion of reconstructions with different foci [19].

Recently, a forward model of image formation in SXT was proposed [24, 25], including rigorous mathematical work [15, 14]. Based on this model, the feasibility and practical application of PSF corrections was shown by Otón et al. [23], where a depth independent correction was applied to SXT data, leading to higher contrast in the obtained reconstruction.

In this manuscript we further generalize the image formation in soft x-ray tomography developed by Otoón et al. [24]. By applying a linear approximation on the model, the effects of a depth dependent PSF can be incorporated into existing iterative reconstruction methods. We present numerical results that show the method is applicable even if the sample is out of focus, or larger than the depth of field. Finally, experimental results show an increase in contrast of a reconstructed image of mouse lymphocytes.

## 2. Image formation in soft x-ray tomography

The understanding of image formation is an important step in any imaging techniques. It allows to choose the best image acquisition strategies and most suitable reconstruction methods. In tomography, the model of image formation has been traditionally based on the Radon transform [29], which is the ideal linear transform (projection) of the specimens attenuation coefficients onto a plane. The projection is linked to the experimental image formation through the Beer-Lambert law, such that the attenuation of the intensity along the ray-paths, *L_i_*, through the sample can be given by

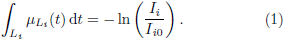

Here *μ_L_i__(t)* is the linear absorption coefficient (LAC) of the specimen along the ray-path, and *I_i_* and *I*_*i*0_ are the attenuated and un-attenuated intensities along the ray-path, respectively.

Although this is a convenient model and a good approximation for highly elongated point spread functions, the image formation in x-ray tomography based on, e.g., diffraction lenses may differ substantially from such ideal model of parallel projections. Despite of this difference, it is beneficial to link the model of image formation in SXT to Eq. (1), so that many available reconstruction methods [21, 12, 9] become available.

Recently, the specifics of the optical system in SXT were integrated into a model of image formation [24, 25]. The model assumes, that propagation of the field, at the vicinity of the sample, can be done by parallel propagation, which is valid if the *NA* of the system is small enough. On the other hand, the model assumes incoherent image formation on the detector, which is reasonable if the *NA* of the condenser and objective lens are matched, so that the image *I_im_* formed at the detector plane *x*_2_ = (*x*_2_,*y*_2_) is given by

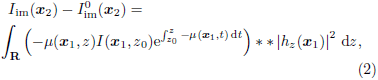

where *I* (*x*_1_, *z*_0_) is the local field intensity at some position *z*_0_ before the sample, **: **R**^2^ → **R**^2^ is the convolution operator in two dimensions, and *h_z_*(***x***_1_) is the impulse response of the optical system. ***I***_im_(***x***_2_) and ***I***^0^_im_(***x***_2_) denotes the recorded images, the latter a reference image without a sample (see Fig. A.7 for details). Depending on the optical setup of the system, these assumptions might not necessarily hold true [31, 20].

Otón et al. [24] describes two cases in which a known solutions exist. In one case, when the impulse response *h_z_*(***x***_1_) is a delta function, the model coincides with Eq. (1). In the other case, the impulse response *h_z_*(***x***_1_) is independent of the axial position *z*. As a result, the ideal Beer-Lambert projections can be recovered by deconvolution of the transmission images, after which conventional reconstruction schemes can be applied.

For a more general solution, in our work, we seek a linear approximation to the forward model Eq. (2), so the image formation in SXT can be expressed as

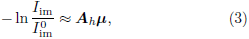

where ***A_h_*** is a linear projection matrix incorporating the PSF of the system, ***μ*** is here a discretized, vector representation of the LAC.

By following the steps of Otoón et al. [24], and building up the left hand side of Eq. (3). The finite difference (see Appendix for details), yields a linear approximation of the form

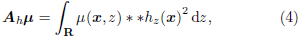

where ***A_h_*** is a projection matrix but now incorporating the depth dependent PSF.

## 3. Numerical results

To validate our, now linear, image formation model, we first performed numerical simulations on a phantom sample, Fig. 2, consisting of binary circles of different sizes.

**Figure 2:**
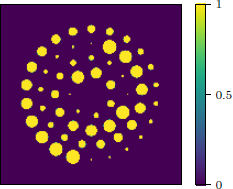
The binary phantom of used for the numerical test. The colormap shown here is scaled with the LAC value of the solid. Fig. 3

Projection images through the phantom sample were calculated using the non-linear model of Eq. (2) on a larger (*L* = 997^1^) discrete grid. The sinograms were down-sampled to size *L* = 256 before tomographic reconstructions. To avoid inversion crimes, the size of high-resolution grid was chosen as a prime number, so that neither the reconstruction or deconvolution was done in the same discrete grid as the calculated phantom.

The theoretical PSF was calculated according to the dimensions of the reconstruction grid and was determined as in Ref. [37], by the converging illumination emerging from a circular lens aperture based on the Huygens-Fresnel principle [2]. The LAC of the phantom was scaled so that the minimum transmission was 0.5*I*_0_.

As a proof of concept, a PSF with a DOF of ± 128, i.e., enclosing the whole sample, and a Rayleigh resolution of 8 units was considered. Two measurements were done, one in-focus case, where the PSF was centered on the center of rotation and one out-of-focus, where the focal spot was shifted to the edge of the image. The sampling criteria of *n* > *L*(π/4) was used for reference and the reconstructions were made using 201 projections^2^. The final projections were distorted by adding Poisson noise.

Three different reconstructions (as shown in Fig. 3) were obtained by solving Eq. (3) by using the Conjugate Gradient Method on the Normal Equations (CGNE): a conventional minimization using the Beer-Lambert approximation, the depth-independent correction of [24], and the linear PSF model.

**Figure 3:**
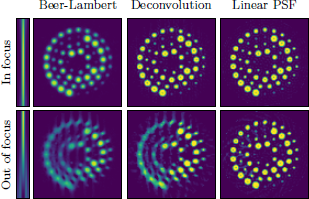
Reconstructions using Beer-Lambert approximation (left) global deconvolution (middle) and PSF projection (right) for two different PSF kernels (far left), both an in-focus PSF (top row) and an out-of-focus PSF (bottom row). Here the images were relatively noiseless with *I*_0_ = 10^6^. Image intensity is the same as for the reference, i.e., the one shown in Fig. 2.

In CGNE, the iteration number can be viewed as a reg-ularization parameter for the solution [8] and running too many iterations will result in amplifying high frequency signals and result in noisy reconstructions. The optimal stopping iterations will of course be data dependent. So as to ensure a fair comparison of the phantom images, we used the oracle knowledge of the phantom to find the highest peak-signal-to-noise-ratio (PSNR)

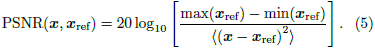

This was done by keeping track of the best solution within the reconstruction scheme and halting when no improvement over the best solution had been recorded within 5 iteration. For the reconstructions shown in Fig. 3 with *I_0_* = 10^6^ the stopping iteration numbers for the Beer-Lambert, deconvolution and PSF reconstructions, respectively were 20,21, and 138 for the in focus PSF, and 20, 21, and 136 for the out of focus PSF.

The optimal deconvolution for the depth-independent correction of [24] was done in two steps – first, noiseless Beer-Lambert projection images were as a convolution kernel to solve for the optimal depth-independent PSF. This 2D kernel was then used to deconvolve the projection images, using the oracle knowledge of the same Beer-Lambert projections as a stopping criteria by minimizing the *l*_2_-norm of the residual.

As seen in Fig. 3, for a centered PSF, the result is as expected. The Beer-Lambert approximation shows smoothing of the edges, characteristic to the PSF. In this case, as the assumption of a depth independent PSF is valid, a proper solution to the inversion exist [24] and global filtering of the projection images yields a good reconstruction result. The linear PSF inversion is similar in quality, but in the case of centered PSF, the depth independent PSF reconstructions have the benefit of much faster convergence.

In the case where the sample is not positioned in the center of PSF, the Beer-Lambert reconstruction clearly shows the artifacts of the defocus. In this case, no suitable global filter exists and the reconstructions based on projection image deconvolution does not improve on the results with respect to BL. Such artifacts are not present in the linear PSF reconstruction.

In numerical simulation, where the projection matrix corresponding to the PSF is known, the linear approximation performs well for the tomographic inversion, as shown in Fig. 3. In practice, however, the image quality is often limited by noise, which makes the inversion problem highly unstable. To investigate in the stability of the inversion, the numerical experiment was repeated using different noise levels. Shown in Fig. 4 the PSNR of the recovered image as a function of the intensity count. As all images were distorted with Poisson noise, a lower count corresponds to a higher noise level. For all cases, the linear PSF inversion provides the highest PSNR and is most resilient to the effects of this noise. For methods when the model of image formation is insufficient, the quality seems to saturate as image quality increases. In this regime, the image model error dominates over the error caused by the noise in the images.

**Figure 4:**
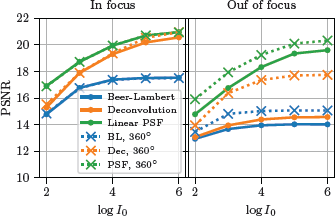
The PSNR Eq. (5) for the reconstructions of the numerical phantom sample as a function of intensity. The solid and dotted lines show 180^°^ and 360^°^ rotation acquisition strategies, respectively.

For out of focus PSF, one can stabilize the reconstructions by taking images over a full, i.e 360°, rotation range. In this fashion, though a sample is not fully in focus at one rotation angle, the sampling at a 180^°^ shift provides additional information, as a different part of the sample will be in focus. Such acquisition can be seen as a modification to the method suggested by Selin et al. [32], where a depth-dependent weight was introduced in the reconstruction scheme to account for projections acquired at different foci. In our case, the projection matrix ***A_h_*** servers a similar function. Such acquisition scheme leads to better reconstruction results in comparison to the 180^°^ rotation, using the same number of projections, with all methods. As seen from the right side image of Fig. 4, the results are even more substantial for the depth independent reconstruction schemes.

## 4. Experimental results

In order to test the method on experimental data, the PSF of the optical system first has to be measured. The measured PSF can then be used to calculate the necessary weights for the projection (and back-projection) operators used in the reconstruction. The experimental work was performed at the National Center for X-ray Tomography at the Advanced Light Source of the Lawrence Berkeley National Laboratory (LBNL). The data was acquired on the XM-2 soft X-ray microscope [18]. The XM-2 is equipped with a cryogenic rotation stage with full 360^°^ range to enable tomographic data collection from cryo-preserved samples. The optical setup consists of two aperture matched Fresnel Zone Plates, with a relatively low NA (0.234), thus the assumptions needed for Eq. (2) should be fairly well met. However, although the theoretical framework rests on the assumption of incoherent image formation, it puts no restrictions into the actual linear operator ***A_h_*** and when measuring the PSF, it is straightforward to incorporate also other effects in the projection matrix, such as various aberrations, distortions, or a spatially varying PSF.

### 4.1. Measuring the system PSF

To measure PSF of the soft x-ray microscope optics, we prepared the phantom sample composed of gold nanoparticles. Spherical gold nanoparticles with diameter of 100 nm (Nanopartz, Cat.No. AR11-100-NB-50) were deposited with a microloader on a 100 nm thick silicone nitride membrane (Silson LtD, Ref.10402101) and then spread by a gentle flow of warm air. The distribution of nanoparticles was confirmed by optical microscopy in dark field mode. The areas with single isolated particles were selected for imaging with x-ray microscope. The regions of interest were imaged with a set of 30 through-focus images with 1 μm step size and 150 ms exposure time each. For each set of radiographs, 20 reference images were acquired with the same exposure time. The radiographs were recorded by a charge-coupled-device camera (Andor IKon-L) with an effective pixel size of 16 nm. A reference profile was obtained by fitting the theoretical intensity function,

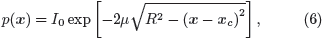

to the experimental data of the in focus image. Here, *I*_0_ is the initial intensity, *μ* is scaled LAC, and ***x_c_*** is center of a sphere.

The PSF was determined by maximum-likelihood (ML) deconvolution [34, 38]. Essentially, we assume that the image of a single bead can be written in form ***y*** = ***p***\*\****h***+***ε***, where the measured signal ***y*** is composed of a convolution between the bead profile, ***p***, the PSF, ***h***, and an additional noise term e. The ML solution for the PSF can now be found iteratively, using the fitted theoretical bead profile Eq. (6), with the Richardson-Lucy algorithm:

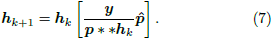

A full 3D PSF was obtained by bilinear interpolation, an example of a PSF extracted in this way is shown in Fig. 5, from which transverse slices give the needed 2D convolution kernels, ***h_z_*** for Eq. (4). The full-width-at-half-maximum of the gaussian peak in focus corresponds to 57 nm, and a maximum loss of intensity of 20% [2, p. 441] along the axial direction gives a DOF of ± 4.7 pm.

**Figure 5:**
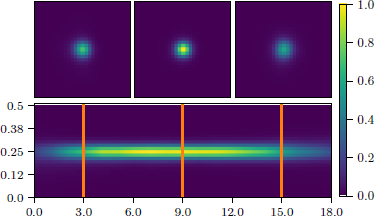
The sagittal slice (along x-ray propagation) ofthe extracted PSF and transverse slices (0.5 μ·0.5 pm) at positions with respect to the focus. The length of the PSF shown here is 18 pm. The shown image intensity is scaled with the maximal value of the PSF.

### 4.2. Experimental validation: imaging of mouse B cell

To test the applicability of the method for experimental data, the method was used to reconstruct tomographic images of mouse lymphocytes. To have full angle rotation during the acquisition, the cells were loaded into thin-wall glass capillaries with a micro loader [26]. For cryo-fixation, the capillaries were rapidly plunged into a liquid propane cooled by liquid N_2_. The data acquisition was done by sequentially rotating the capillary with 2° increments for 180° with an exposure time of 300 ms per image. A series of 10 reference images were taken before and after the scan, that were used to normalize the data. The projection images were aligned using an updated version of the previously developed automatic registration software AREC3D [27].

The tomographic image was reconstructed in three different ways using CGNE, viz.:, A reference image using the Beer-Lambert approximation, a reconstruction where the intensity images are deconvolved with a 2D PSF, i.e., the solution to Eq. (2) if the PSF is not depth-dependent, and using the linear PSF approximation presented in this paper. The 2D kernel for the deconvolution was obtained by taking the average of all the measured 2D kernels within the DOF of the PSF. The deconvolution was performed on the transmission images using RL deconvolution, and stopped before significant ringing was observed^3^.

In Fig. 6 we show details of the reconstructions showing endoplasmic reticulum for the three different reconstructions. The linear PSF approximation performs as well as the deconvolution method, which can, in this case, be considered to be a proper solution to the inversion as the DOF of the PSF spans the whole sample. Both PSF corrected reconstructions show an increase of contrast with respect to the Beer-Lambert approximation. The effect is less pronounced in the in-plane reconstruction, where image quality is limited by the relatively sparse sampling.

**Figure 6:**
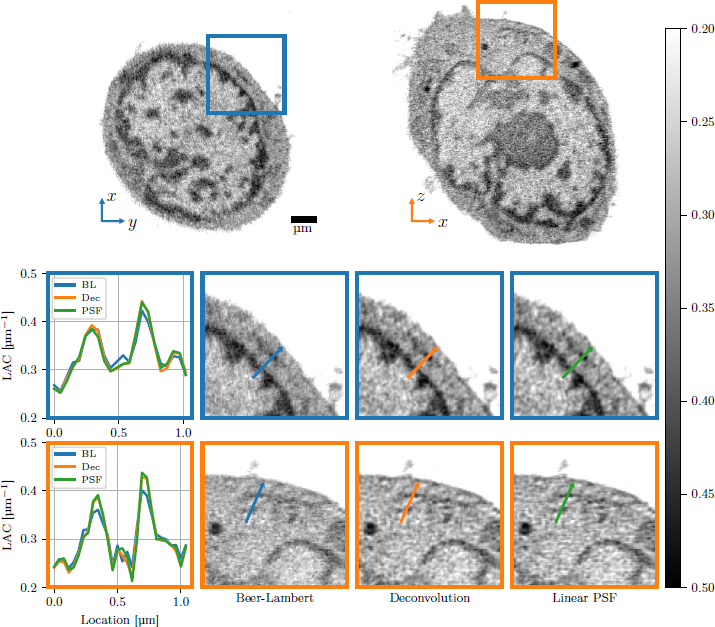
An in-plane slice (plane orthogonal to the axis of rotation) and a transverse plane of the reconstruction of a mouse B-cell. The two rows show regions of interest (3.5 μ·3.5 μ·) from the corresponding slice. The arrows correspond to line profile depicted in the corresponding leftmost plot, which shows the LAC profile along the line.

## 5. Conclusions

In this work we have introduced a linear approximation to the reconstruction of tomographic x-ray data, including the effect of the point spread function of the optics. We show numerically, that the approximation is well suited for the inversion of the SXT data both when the PSF acts as depth-independent blur as well as when the sample is partially out of focus.

For experimental measurements, the PSF inversion increases contrast in the image, especially of edge features, and provides similar results as compared to deconvolution. Numerical results show that the introduced inversion using a projection matrix including the PSF is more resilient to noise than its deconvolution-based counterpart. This, however, comes at a price, since the reconstruction using the PSF incorporated projection matrix is both computationally more expensive, and converges significantly slower.

The presented model tackles current limitations of SXT and provides new ways of data acquisition. For a properly sampled and focused PSF, the linear PSF approximation enables use of higher NA optics, as the limited depth of field can be computationally amended. The results in Fig. 4 predict a possible way to extend the depth of field experimentally by shifting the focus to either side of the sample and taking a full 360^°^ rotation data set. This would allow one to circumvent the traditional limitations of diffraction limited optics Fig. 1 and extend the effective depth of field with respect to the resolution. In other words, it would either enable imaging current samples with a higher NA, thus a higher resolution, or extending the DOF of current imaging systems, allowing for full 3D reconstructions of larger samples.

We expect the method to be generally applicable also for other tomographic systems, as log as the assumption of a linear image formation can be met.

## Acknowledgment

Research reported in this publication was conducted at The National Center for X-ray Tomography, located at the Advanced Light Source of Lawrence Berkeley National Laboratory, and was supported by grants from NIH (P41GM103445), DOE's Office of Biological and Environmental Research (DE-AC02-5CH11231), and Chan Zuckerberg Initiative Human Cell Atlas program. Venera Weinhardt was funded by German Research Foundation fellowship WE6221/1-1.

## Appendix A Linear approximation of the image formation

The model of Otón et al. [24] is based on two assumptions, in which the measured projection is a result of the attenuated light passing through the sample, blurred by the PSF of the objective lens. We assume firstly, that there exists an impulse response function ***h_z_***(***x***_1_, ***x***_2_) (see e.g. Refs [2] and [7] for details), such that the intensity field at the image plane, ***I***_im_(***x***_2_), can be expressed by linear transport of an (un-attenuated) field intensity *U* at position *z*,

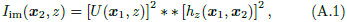

where we define ***x***_1_ and ***x***_2_ as two sets of coordinates, corresponding to the planes perpendicular to the optical axis at *z*_1_ and *z*_2_ and **: **R**^2^ → **R**^2^ is the convolution operator in two dimensions. The second assumption is, that the local field propagation within teh sample can be done by parallel wave approximation.

**Figure A.7:**
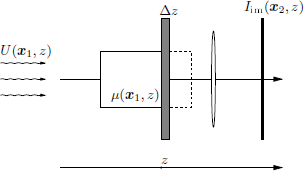
The PSF projection is constructed by assuming the image of a “partially cut” sample can be incoherently transferred to the image plane by linear transport. The resulting image *I* of the whole sample can be constructed by finite difference by adding slices of thickness d*z*.

For cleaner notation, we construct *h_z_*(***x***_1_, ***x***_2_) so that it includes all linear scaling (magnification, inversion), thus we can express the field transfer simply as a 2D convolution of the local field plane and a 2D PSF [*h_z_* (***x***)]^2^. Using these assumptions Otoón et al. [24] construct the derivative of *I*_im_*^z^* with respect to *z* by considering a partially cut sample and adding to it slices of thickness Δ*z* (see Fig. A.7).

Following the derivation by Otoón et al. [24], we construct the image formation by finite difference of the image, but applying this directly on the logarithm of the normalized image, that is we seek an approximation to the normalized image

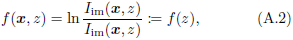

where we drop the explicit notation of ***x*** for cleaner notation. Constructing the finite difference and substituting for the linear transfer in Eq. (A.1) yields

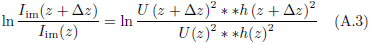

For the left hand side, we note that from the series expansion

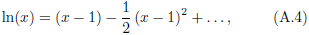

we can neglect higher order terms as, *x* ≈ 1. For the right hand side, making the same approximation as Otóon et al. [24] we propagate the field coherently (to linear accuracy)

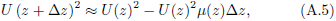

and as the PSF can be assumed smooth, we can also assume that

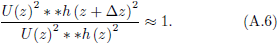

yielding

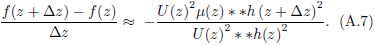

To get rid of the final non-linear terms, we assume that *U(z)*^2^ is smooth enough, such that

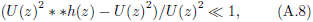

yielding

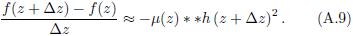

Integrating on both sides of this finite difference approximation of the derivative, we get

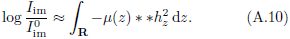

With suitable discretization, we can now express the right hand side as a linear projection matrix ***A_h_μ***, yielding our approximation, Eq. (3).

### Appendix A.1. Validity of the linear approximation

To test the assumptions for the forward model, we calculated the difference on the three forward models of a reconstruction of a biological sample taken with XM2 on beamline 2.1 at the ALS in LBNL [18]. In Fig. A.8 we show the error of a Beer-Lambert projection and our linear PSF approximation, respectively, as compared to the non-linear projection model of Eq. (2). It is evident that the error of the linear approximation is negligible except for at the edges of the samples. This is expected, as the samples in XM2 are mounted in capillary tubes. The capillary tubes produce large gradients in the LAC, where the assumption of a relatively smooth local field in Eq. (A.8) is expected to fail.

**Figure A.8:**
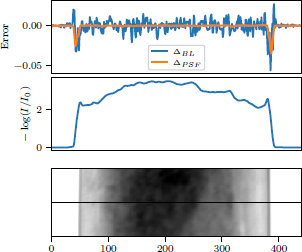
The error of the linear model of the projection as compared to the non-linear model of Eq. (2). Δ*_BL_* and Δ*_PSF_* are to the errors of the linear approximation using parallel projection, log(*I*/*I*_0_) + ***Ax***, and PSF projection, log(*I*/*I*_0_) + ***A***_*h*_*x*, respectively.

## Footnotes

1 All lengths describing the numerical validation, are in units of pixels.

2 The odd number of projections is convenient, as it ensures that the same sampling could be used for both 180 and 360°. If *n* is even, and the sampling is done over 360°, the angles will be at a π shift and do not provide any additional sampling if the PSF is in focus and symmetric with respect to the defocus.

3 Of course, as the experimental data lacks oracle information this could not be done in as rigorous manner as for the phantom sample, so the “best” result had to be qualitatively determined by visual inspection.

## References

[1] Attwood D. (2007). Soft x-rays and extreme ultraviolet radiation: principles and applications. Cambridge University Press.

[2] Born M. and E. Wolf (1970). Principles of Optics: Electromagnetic Theory of Propagation, Interference and Diffraction of Light. Pergamon Press Ltd, Headington Hill Hall, Oxford.

[3] Carzaniga R., M.-C. Domart, L. M. Collinson, and E. Duke (2013, nov). Cryo-soft X-ray tomography: a journey into the world of the native-state cell. Protoplasma 251 (2), 449–458.

[4] Chen H.-Y., D. M.-L. Chiang, Z.-J. Lin, C.-C. Hsieh, G.-C. Yin, I.-C. Weng, P. Guttmann, S. Werner, K. Henzler, G. Schneider, et al. (2016). Nanoimaging granule dynamics and subcellular structures in activated mast cells using soft x-ray tomography. Scientific reports 6, 34879.

[5] Chiappi M., J. J. Conesa, E. Pereiro, C. O. S. Sorzano, M. J. Rodriguez, K. Henzler, G. Schneider, F. J. Chichon, and J. L. Carrascosa (2016, mar). Cryo-soft X-ray tomography as a quantitative three-dimensional tool to model nanoparticle:cell interaction. Journal of Nanobiotechnology 14 (1).

[6] Duke E., K. Dent, M. Razi, and L. M. Collinson (2014, jun). Biological applications of cryo-soft X-ray tomography. Journal of Microscopy, 65–70.

[7] Goodman J. W. (1996). Introduction to Fourier optics (Second ed.). THE McGRAW-HILL COMPANIES, INC. New York St. Louis San Francisco Auckland Bogot6 Caracas Lisbon London Madrid Mexico City Milan Montreal New Delhi San Juan Singapore Sydney Tokyo Toronto.

[8] Hansen P. C. (1998). Rank-deficient and discrete ill-posed problems: numerical aspects of linear inversion, Volume 4. Siam.

[9] Herman G. T. (2009). Fundamentals of computerized tomography: image reconstruction from projections. Springer Science & Business Media.

[10] Hertz H., O. von Hofsten, M. Bertilson, U. Vogt, A. Holmberg, J. Reinspach, D. Martz, M. Selin, A. Christakou, J. Jerlström-Hultqvist, and S. Svard (2012, feb). Laboratory cryo soft X-ray microscopy. Journal of Structural Biology 177(2), 267–272.

[11] Jensen G. J. and R. D. Kornberg (2000, jul). Defocus-gradient corrected back-projection. Ultramicroscopy 84 (1-2), 57–64.

[12] Kak A. C. and M. Slaney (1988). Principles of computerized tomographic imaging. IEEE press.

[13] Kazantsev I. G., J. Klukowska, G. T. Herman, and L. Cernetic (2010). Fully three-dimensional defocus-gradient corrected back-projection in cryoelectron microscopy. Ultramicroscopy 110(9), 1128–1142.

[14] Klukowska J. and G. T. Herman (2014). Reconstruction from Microscopic Projections with Defocus-Gradient and Attenuation Effects, pp. 157–186. New York, NY: Springer New York.

[15] Klukowska J., G. T. Herman, J. Oton, R. Marabini, and J.-M. Carazo (2014, dec). The soft x-ray transform. Inverse Problems 30(12), 125015.

[16] Kohl H. and L. Reimer (2008). Transmission Electron Microscopy: Physics of Image Formation (5 ed.). 1556–1534. Springer New York.

[17] Le Gros, M. A., E. J. Clowney, A. Magklara, A. Yen, E. Markenscoff-Papadimitriou, B. Colquitt, M. Myllys, M. Kellis, S. Lomvardas, and C. A. Larabell (2016). Soft x-ray tomography reveals gradual chromatin compaction and reorganization during neurogenesis in vivo. Cell reports 17(8), 2125–2136.

[18] Le Gros M. A., G. McDermott, B. P. Cinquin, E. A. Smith, M. Do, W. L. Chao, P. P. Naulleau, and C. A. Larabell (2014). Biological soft X-ray tomography on beamline 2.1 at the Advanced Light Source. Journal of synchrotron radiation 21 (6), 1370–1377.

[19] Li F., Y. Guan, Y. Xiong, X. Zhang, G. Liu, and Y. Tian (2017). Method for extending the depth of focus in x-ray microscopy. Optics Express 25 (7), 7657–7667.

[20] McNally J. G., S. Rehbein, C. Pratsch, S. Werner, G. Schneider, et al. (2016). 3D PSF Measurement for a Soft X-ray Microscope and Comparison to Theory. In Computational Optical Sensing and Imaging. Optical Society of America.

[21] Natterer F. (1986). Computerized Tomography, pp. 1–8. Wiesbaden: Vieweg+Teubner Verlag.

[22] Otón J., E. Pereiro, J. J. Conesa, F. J. Chichón, D. Luque, J. M. Rodriguez, A. J. Perez-Bernó, C. O. S. Sorzano, J. Klukowska, G. T. Herman, et al. (2017). Xtend: Extending the depth of field in cryo soft x-ray tomography. Scientific Reports 7.

[23] Otoón J., E. Pereiro, A. J. Póerez-Bernóa, L. Millach, C. O. S. Sorzano, R. Marabini, and J. M. Carazo (2016). Characterization of transfer function, resolution and depth of field of a soft x-ray microscope applied to tomography enhancement by wiener deconvolution. Biomedical Optics Express 7(12), 5092–5103.

[24] Otoón J., C. Sorzano, E. Pereiro, J. Cuenca-Alba, R. Navarro, J. M. Carazo, and R. Marabini (2012, apr). Image formation in cellular X-ray microscopy. Journal of Structural Biology 178(1), 29–37.

[25] Otóon J., C. O. S. Sorzano, F. J. Chichóon, J. L. Carrascosa, J. M. Carazo, and R. Marabini (2014). Soft X-Ray Tomography Imaging for Biological Samples, pp. 187–220. New York, NY: Springer New York.

[26] Parkinson D. Y., L. R. Epperly, G. McDermott, M. A. Le Gros, R. M. Boudreau, and C. A. Larabell (2013). Nanoimaging cells using soft X-ray tomography. Nanoimaging: Methods and Protocols, 457–481.

[27] Parkinson D. Y., C. Knoechel, C. Yang, C. A. Larabell, and M. A. Le Gros (2012). Automatic alignment and reconstruction of images for soft X-ray tomography. Journal of structural biology 177(2), 259–266.

[28] Patwardhan A., A. Ashton, R. Brandt, S. Butcher, R. Carzaniga, W. Chiu, L. Collinson, P. Doux, E. Duke, M. H. Ellisman, E. Franken, K. Gruönewald, J.-K. Heriche, A. Koster, W. Kuöhlbrandt, I. Lagerstedt, C. Larabell, C. L. Lawson, H. R. Saibil, E. Sanz-Garcóia, S. Subramaniam, P. Verkade, J. R. Swedlow, and G. J. Kleywegt (2014, oct). A 3D cellular context for the macromolecular world. Nature Structural & Molecular Biology 21 (10), 841–845.

[29] Radon J. (1917). On determination of functions by their integral values along certain multiplicities. Berichte der Săchische Akademie der Wissenschaften Leipzig,(Germany) 69, 262–277.

[30] Schneider G., P. Guttmann, S. Heim, S. Rehbein, F. Mueller, K. Nagashima, J. B. Heymann, W. G. Muöller, and J. G. McNally (2010, nov). Three-dimensional cellular ultrastructure resolved by X-ray microscopy. Nature Methods 7(12), 985–987.

[31] Selin M., E. Fogelqvist, A. Holmberg, P. Guttmann, U. Vogt, and H. M. Hertz (2014). 3D simulation of the image formation in soft x-ray microscopes. Optics Express 22(25), 30756–30768.

[32] Selin M., E. Fogelqvist, S. Werner, and H. M. Hertz (2015). Tomographic reconstruction in soft x-ray microscopy using focus-stack back-projection. Optics letters 40(10), 2201–2204.

[33] Uchida M., Y. Sun, G. McDermott, C. Knoechel, M. A. L. Gros, D. Parkinson, D. G. Drubin, and C. A. Larabell (2010, dec). Quantitative analysis of yeast internal architecture using soft X-ray tomography. Yeast 28(3), 227–236.

[34] van Kempen G. M., H. T. van der Voort, J. G. Bauman, and K. C. Strasters (1996). Comparing maximum likelihood estimation and constrained tikhonov-miller restoration. IEEE Engineering in Medicine and Biology Magazine 15(1), 76–83.

[35] Voortman L. M., E. M. Franken, L. J. van Vliet, and B. Rieger (2012). Fast, spatially varying ctf correction in tem. Ultrami-croscopy, 118 26–34.

[36] Voortman L. M., S. Stallinga, R. H. Schoenmakers, L. J. van Vliet, and B. Rieger (2011, jul). A fast algorithm for computing and correcting the CTF for tilted, thick specimens in TEM. Ultramicroscopy 111 (8), 1029–1036.

[37] Weiß D., G. Schneider, B. Niemann, P. Guttmann, D. Rudolph, and G. Schmahl (2000). Computed tomography of cryogenic biological specimens based on X-ray microscopic images. Ultramicroscopy 84(3–4), 185–197.

[38] Yoo H., I. Song, and D.-G. Gweon (2006). Measurement and restoration of the point spread function of fluorescence confocal microscopy. Journal of microscopy 221 (3), 172–176.

